# Drosben, an affordable system for scalable survival analysis in *Drosophila*

**DOI:** 10.64898/2026.07.02.736118

**Authors:** Terrence M. Trinca, Pau Berenguer-Molins, Cristina Fernández-García, Joaquín de Navascués

**Affiliations:** European Cancer Stem Cell Research Institute, School of Biosciences, Cardiff University, Cardiff, United Kingdom; School of Biochemistry, University of Bristol, Bristol, United Kingdom; Hospital del Mar Research Institute (HMRI), Barcelona, Spain; School of Life Sciences, University of Essex, Colchester, United Kingdom

**Keywords:** *Drosophila*, survival, lifespan, ageing, open hardware, magnetic fields, auxin

## Abstract

Survival analysis is a workhorse assay in *Drosophila* research to evaluate somatic fitness. It is indispensable in the study of ageing and insightful in immunity, metabolism, radiobiology, toxicology, ecology, and others. While conceptually simple, lifespan measurement is labour-intensive because it requires the continuous manual maintenance of large experimental cohorts. Here, we describe Drosben, an approach that combines a 3D-printed device to transfer flies from several vials simultaneously, a paper system for quick data recording and accompanying software that automatically digitalises life tables for analysis. We show that using Drosben reduces the time investment to perform lifespan assays by ∼85% respect to handling individual vials, with improved speed regardless of user experience handling these. Using Drosben, we address the effects on longevity of chronic feeding of indole-acetic acid (IAA), naphthalene-acetic acid (NAA) and trimethoprim (TMP) – compounds used to control heterologous targeted protein degradation systems. We find that IAA and NAA mildly shorten lifespan, while TMP extends female longevity. We further show that strong static magnetic fields do not affect *Drosophila* lifespan. Our work suggests that Drosben can cheaply accelerate research where lifespan is used as a life history trait.

## INTRODUCTION

For over a century, *Drosophila melanogaster* has contributed to landmark discoveries in developmental biology, cancer, immunity, neurobiology, ageing, genetics and genomics (Rubin and Lewis 2000, Helfand and Rogina 2003, Lemaitre 2004, Letsou and Bohmann 2005, Bellen, Tong et al. 2010, Gonzalez 2013, Villegas 2019). This laid the foundation to exploit the fruit fly for studying an ever-growing number of human diseases, from their basic biology to drug screening (Whitworth, Wes et al. 2006, Bharucha 2009, Johnson and Cagan 2010, Rand 2010, Pandey and Nichols 2011, Willoughby, Schlosser et al. 2013, Ugur, Chen et al. 2016, Wangler, Yamamoto et al. 2017, Musselman and Kühnlein 2018, Oriel and Lasko 2018, Staats, Lüersen et al. 2018, Papanikolopoulou, Roussou et al. 2019, Demir 2020, Trinca and de Navascues 2025).

Survival analyses provide fundamental information about the influence of genetics and the environment on overall organismal health (Mair, Goymer et al. 2003, Flatt 2011, Tatar, Post et al. 2014, Galenza, Hutchinson et al. 2016, Strilbytska, Semaniuk et al. 2020). Moreover, low culturing costs and short lifespan of *Drosophila* (typically under 100 days) allows performing well-powered lifespan experiments, with hundreds to thousands of experimental individuals. This makes survival analysis a staple experiment in many *Drosophila* laboratories.

However, *Drosophila* lifespan analyses are labour-intensive as they require frequent ‘flipping’: the passaging of survivors into fresh medium by manual inversion of old vials and gentle banging of the flies into fresh tubes, individually (Linford, Bilgir et al. 2013). The recording of deaths is an additional, time-consuming activity. The manual mechanics of flipping and record keeping limit the scalability of maintaining concurrently running survival experiments. While there are specialised robots that can flip tubes automatically (Stowers Institute for Medical 2011), these are only cost-effective for large facilities, where they offset the extraordinary effort to care for thousands of strains (Bangham 2019). Moreover, current automated systems are not appropriate for lifespan measurement, as they typically use anaesthetics during transfer and do not record events. The commercially available ‘Drosoflipper’ device has been developed recently to reduce the time of manual handling (https://www.drosoflipper.com); however, it does not address the time-consuming inefficiencies associated with data recording.

The availability of affordable electronic components, design software and 3D-printing has enabled the development of home-made, bespoke lab equipment at unprecedented scale, from basic micropipettes to microfluidic chambers or fluorescent microscopes (Baden, Maia Chagas et al. 2015, Yazdi, Popma et al. 2016, Maia Chagas, Prieto-Godino et al. 2017, Machado, Malpica et al. 2019, Trinca, Peña et al. 2025). We took advantage of the current affordability of 3D-printing to develop an open-source system for increased throughput in *Drosophila* lifespan experiments, which we describe here. In *Drosophila* tradition, we named this system *Drosben*, a portmanteau of “*Drosophila*” and “trosben”—Welsh for somersault. Drosben comprises pieces of hardware and accompanying software. The hardware components are easily assembled out of 3D-printed parts and a few off-the-shelf parts. The software generates printable experiment-specific data-recording forms for use at the bench, compiles the data automatically, and produces a preliminary analysis. Compared to passaging vials individually (henceforth, ‘conventional’ handling), using Drosben reduced handling time ∼5-fold in both inexperienced and experienced drosophilists. As the system includes magnets and it has been claimed that *Drosophila* can sense static magnetic fields and have their behaviour affected by artificial ones (Gegear, Casselman et al. 2008, Fedele, Green et al. 2014), we used Drosben to test whether strong magnetic fields affect *Drosophila* longevity, and found no significant effects. Next we considered the growing usage of small molecule-regulated degron systems like the Auxin Inducible Degron (AID) and the ecDHFR Destabilizing Domain (DD) for regulating transgene expression *Drosophila* (Trost, Blattner et al. 2016, Bence, Jankovics et al. 2017, Sethi and Wang 2017, Chen, Werdann et al. 2018, Kogenaru and Isalan 2018, Lopez Del Amo, Leger et al. 2020, Jullien, Guillou et al. 2022, McClure, Hassan et al. 2022), and the potential effects of chronic feeding of these molecules on lifespan. Using Drosben, we tested two auxins, indole-3-acetic acid (IAA) and the synthetic 1-naphthalene-acetic acid (NAA), as well as the DD stabiliser, the antibiotic trimethoprim (TMP). We found that, at concentrations used for regulating their corresponding degrons, TMP prolongs female lifespan while IAA and NAA slightly reduce survival in both sexes.

## MATERIALS AND METHODS

### *Drosophila* strains and husbandry

All stocks were maintained at 25°C with a 12 h:12 h light:dark photocycle. Food was made from organic yellow maize flour (80 g/L), inactivated yeast powder (30 g/L), Brewer’s dextrose (80 g/L), agar (6.67 g/L), cooked at 95°C before adding propionic acid (0.5%) and tegosept (0.005%). Flies were housed in polystyrene narrow vials (P1068-T, Regina Scientific Supplies, Newcastle-under-Lyme, UK) with dimensions 95 mm high × 25 mm outer diameter and containing 3-10 mL of food, depending on the experiment. We used strains *w^1118^* (RRID:BDSC_5905) and *Vallecas* (Morata and Garcia-Bellido 1973) (a gift from Prof Sonsoles Campuzano).

### 3D modelling and printing

3D modelling was performed in FreeCAD v1.0.2 (Riegel, Mayer et al. 2016) and Autodesk Fusion 2703.1.20 (Autodesk Inc., San Francisco, USA). Models are available in GitHub (https://github.com/jdenavascues/Drosben) and preserved permanently in Zenodo (Trinca and de Navascués 2026) in both editable (FCSts) and printable (3mf) non-proprietary file formats. All components were benchmarked for printing on an i3 Mega (Anycubic) and further tested for printing on models i3 MK3S+ (Prusa) and A1mini (Bambulab). Models printed well in either polyactic acid (PLA) or acrylonitrile butadiene styrene (ABS) with 1.75 mm filaments and using 0.4 mm nozzles. While ABS is more thermotolerant than PLA, it is also more expensive; we did not find signs of wear or mechanical weakening in PLA components after months in *Drosophila* incubators at 25-30 °C, so we did most of our printing in PLA. Slicing was done with either UltiMaker (Cura) or BambuStudio (Bambulab). None of the designs required specific slicing parameters; we used 25% infill for strength, supports where necessary, and layer height < 0.2 mm.

### Hardware assembly

Most parts required removing printed supports using pliers. Some parts were not functional directly as printed and required gluing additional components. We outline here the essential information; additional details are available at the Drosben online manual https://jdenavascues.github.io/Drosben.

#### Rack and lid

An N52 Neodymium-Iron-Boron disc (12 mm radius × 3 mm thickness) magnet (Hhoomy Magnet Co. Ltd.) was placed into the hole on top of the rack using cyanoacrylate gel glue.

#### Multiflipper

Twelve ∼90 mm-long tubes of extruded acrylic (25 mm outer diameter, 2 mm thickness) (Simply Plastics, Essex, UK) were glued to each other with acrylic cement (Tensol 12) and the group to the flat side of the Multiflipper body (with cyanoacrylate). Tubes were covered on the opposite side with glued cloth: either nylon mesh or woven stainless steel wire mesh (200 lines per inch, 0.07 mm hole size, 0.05 mm wire thickness) (The Mesh Company, Warrington, UK).

#### Depositor, lid and slider

For rigidity, 40 mm carbon steel nails (with the heads cut off with pliers) were inserted at both ends of the hollow rails of the depositor and lid, glued with cyanoacrylate gel. Another one was inserted and glued in the cylindrical cavity at the far end of the slider, so it would be pulled by the magnets in the multiflipper. A piece of 1 mm thick neoprene rubber sheet (Camthorne Industrial Supplies, Stoke-on-Trent, UK) was cut using the 3D-printed depositor stencil and glued to the depositor hollow outline. Cross cuts were done where the rubber covers the holes of the depositor to make a cuspid valve in each. Non-printed parts and materials were bought from various online retailers in the UK, but widely available internationally (**Table 1**).

**Table 1:** Bill of materials to manufacture the Drosben components. List of materials, recipes and usage requirements to make a cost behaviour analysis to scale Drosben use.

| Component costs |  |  |  |  |
| --- | --- | --- | --- | --- |
| Component | Quantity | Required <sup>(a)</sup> | Cost per usage unit |  |
| (initial materials) | 1 | Per lab | £71.00 |  |
| Depositor stencil | 1 | Per lab | £0.14 |  |
| Multiflipper | 1 | Per user | £38.91 |  |
| Slider | 1 | Per user | £0.50 |  |
| Depositor | 1 | Per user | £3.49 |  |
| Rack | 2 | Per cohort | £2.28 |  |
| Lid | 1 | Per cohort | £0.90 |  |
| Materials costs |  |  |  |  |
| Material | Quantity (per pack) | Unit | Price <sup>(b)</sup> |  |
| PLA filament | 1000 | g | £18.00 |  |
| Acrylic tube | 1 | count | £3.00 |  |
| Nail shaft | 200 | count | £3.50 |  |
| N52 magnet 12 x 3 mm | 50 | count | £12.00 |  |
| N52 magnet 10 x 3 mm | 50 | count | £12.00 |  |
| Woven wire mesh | 3600 | cm<sup>2</sup> | £20.00 |  |
| Super glue gel | 30 | g | £4.00 |  |
| Acrylic cement | 50 | ml | £9.00 |  |
| Rubber sheet | 100 | cm<sup>2</sup> | £3.00 |  |
| Recipes |  |  |  |  |
| Component | Material | Quantity | Unit | Cost per component |
| Multiflipper | PLA filament | 40 | g | £0.72 |
|  | Acrylic tube | 12 | count | £36.00 |
|  | N52 magnets 10 x 3 mm | 4 | count | £0.96 |
|  | Woven wire mesh | 120 | cm<sup>2</sup> | £0.67 |
|  | Super glue gel | 0.2 | g | £0.03 |
|  | Acrylic cement | 3 | ml | £0.54 |
| Depositor | PLA filament | 21 | g | £0.38 |
|  | Rubber sheet | 100 | cm<sup>2</sup> | £3.00 |
|  | Nail shaft | 4 | count | £0.07 |
|  | Super glue gel | 0.3 | g | £0.04 |
| Lid | PLA filament | 32 | g | £0.58 |
|  | N52 magnet 12 x 3 mm | 1 | count | £0.24 |
|  | Nail shaft | 4 | count | £0.07 |
|  | Super glue gel | 0.1 | g | £0.01 |
| Rack | PLA filament | 50 | g | £0.90 |
|  | N52 magnet 12 x 3 mm | 1 | count | £0.24 |
| Slider | PLA filament | 26 | g | £0.47 |
|  | Nail shaft | 1 | count | £0.02 |
|  | Super glue gel | 0.1 | g | £0.01 |
| Depositor stencil | PLA filament | 8 | g | £0.14 |

Tools
| *Tool* | *Assumed available* | *Cost* |
| --- | --- | --- |
| Pliers (to clip nail heads) | yes | £10.00 |
| Cutter (to cut mesh and rubber sheet) | yes | £10.00 |
| 3D printer | yes | £140.00 |
| Silicon sheet, 0.5 mm thick (to glue the multiflipper) | no | £4.50 |

### Accompanying software

The software part of Drosben was written in Python 3.11 (Python Software Foundation) using Jupyterlab 4.5.5 (Jupyter Foundation) and R 4.6.0 (Team 2026) using RStudio 2026.05.1+225 (Posit Software, PBC). The full Python dependencies are recorded in the GitHub repository ‘environment-pinned.yml’ file. R code was mostly based on packages *survival* (Therneau 2026) and *survminer* (Kassambara, Kosinski et al. 2026).

Instructions to install the Drosben software are available in the GitHub repository (https://github.com/jdenavascues/Drosben) and the Drosben online manual. The steps described in the manual are implemented as Jupyter (Granger and Pérez 2021) and R markdown (Allaire, Xie et al. 2026) notebooks available in GitHub, that the user can save, modify and run locally.

### Scanning

Drosben requires scanning at two points: to ensure that the colour markers used to record deaths, carried-overs and censorings can be distinguished automatically (colour calibration step), and to extract and digitally log the data from the paper datasheets. The most important consideration for this is to use the same hardware for scanning both the colour calibration sheets and the datasheets. The second consideration is resolution: colour calibration sheets need to be scanned at ≥150 pixels per inch; datasheets need ≥200, and be in good orientation (the software can correct the angle but only up to a few degrees). The final consideration is the hardware. We found flatbed scanners to give the best results: they are slow but provide homogeneous, reproducible illumination, no perspective distortion, and custom resolution. Photocopy machines are faster and provide good illumination and orientation, but do not allow choice of resolution. Phones will also work, provided you can ensure homogeneous, reproducible illumination (e.g. with an LED ring), frontal imaging (e.g. with a rig) and papers with no creases or folds (e.g. recently used or flattened with non-reflective clear sheet).

### Measuring handling time with volunteer participants

Eleven participants recruited from the Cardiff University *Drosophila* research community were asked to transfer a cohort of 120 *w^1118^*females in 12 vials using conventional flipping and Drosben. We recorded the time taken to transfer flies with each method (once for the conventional, as it aggregates the individual flipping of 12 tubes each, and three times for Drosben, taking the average) and their experience level handling *Drosophila*. Ethical approval to record performance of human volunteers was granted by the School of Biosciences (Cardiff University) Research Ethics Committee (SREC 1811-02).

### Lifespan assays

All cohorts were age synchronised as described previously (Clancy and Kennington 2001, Linford, Bilgir et al. 2013, Piper and Partridge 2016). Upon adult emergence, flies were transferred to fresh food to mate for 48 hours. Flies were then sorted by sex, deposited in the Drosben housing racks (10-15 flies per vial) and maintained using the Drosben system (**Figure 1** and **Supplementary Figure 1**). *Drosophila* were passaged to fresh food 3 times per week, with dead, escaped (censored) or carried over (dead bodies falling into fresh tubes) recorded at each transfer using the Drosben datasheets (**Figure 1N-O**). The Drosben software was used to generate experiment-specific data logging forms (**Supplementary Figure 2**) prior to the experiment and to automatically compile the data thereafter.

**Figure 1.**
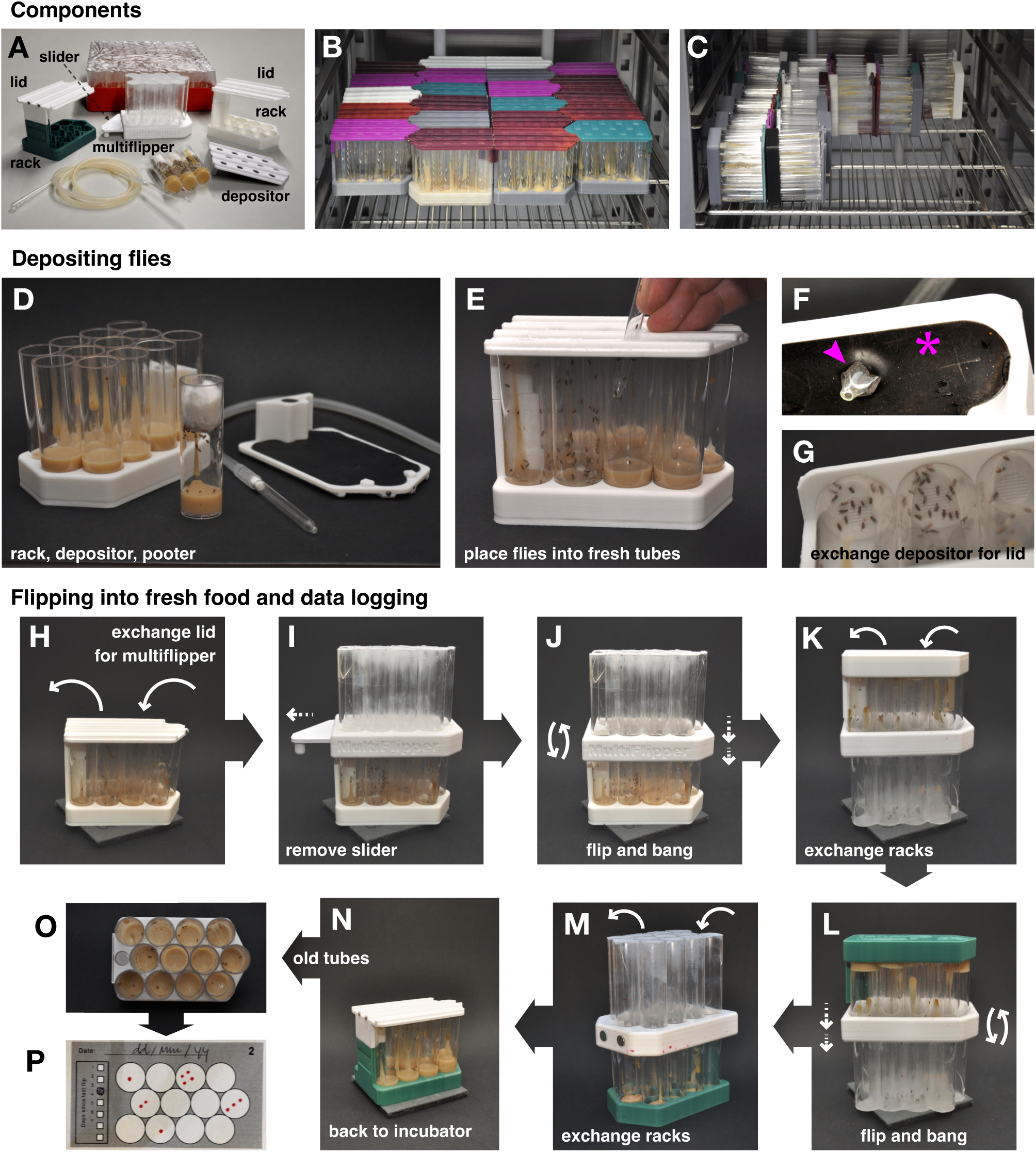
Components and usage of the Drosben system. (**A**) Components needed for initiating (depositor, rack, lid, food vials and pooter) and passaging (multiflipper, slider, second rack) an experiment. (**B-C**) Racks can be arranged compactly in incubator shelves. When laying on their base, 24 racks fit in a 50 cm wide × 54 cm deep shelf (**B**, only 22 racks shown). When laying on their spine, up to 35 racks fit in the same area (**C**, only 22 are shown). (**D-G**) Steps for depositing flies. Fresh, racked vials (**D**) are covered with the depositor (**E**), and flies are blown into each vial with the pooter, across the cuspid valves. (**F**) shows a traversed rubber valve (arrowhead) and a closed one (asterisk). The depositor is then exchanged for the lid, which secures the flies long-term (**G**). (**H-P**) Steps for flipping and recording observations. The lid of the rack with old vials (**H**) is replaced with the multiflipper, whose chambers are closed by the slider (**I**). The slider is then removed, connecting each tube in the rack with a chamber (**J**). The assembly is flipped vertically, and flies are gently banged into the multiflipper (**K**). The old rack is quickly exchanged for one with fresh food (**L**) and set aside. Multiflipper and new rack are flipped and flies banged into the rack (**M**). The multiflipper is exchanged for a lid (**N**), the new rack goes back to the incubator (**B-C**). The old tubes are inspected for dead stuck flies (**O**), which are recorded in the corresponding datasheet (**P**).

Indole-3-acetic acid (Merck I2886), 1-naphthaleneacetic acid (Merck N0640) and trimethoprim (Merck 92131) were supplemented into *Drosophila* food just prior to pouring into vials. Auxins were dissolved in ethanol and trimethoprim in dimethyl sulfoxide (50 mg/ml) with control sham food prepared with equal volumes of solvent. *Drosophila* were then maintained on these vials as an *ad libitum* diet for the duration of their lifespan and passaged using Drosben or conventional flipping.

### Survival statistics

Time-to-event statistical analysis was performed in R (see Accompanying software) using *p*<0.05 as threshold for significance in all tests. We used the log-rank test for trend to determine whether magnetic field affected longevity, as the magnitude of the effect was not of interest. We used Cox’ regression for all other experiments, after verifying proportional hazards (PH) with Schoenfeld tests, so we could estimate the magnitude of effects. These are reported as hazard ratios (HR) with their corresponding *p*-value from a Wald’s test. For multiple-variable experiments, we used Cox’ regression to model both the additive and interactive effects of the variables and report, for the best model, the HR and Wald’s test *p*-value for each of the individual variables and, when relevant, their interaction term.

## RESULTS

### Overview of the Drosben approach to lifespan analysis

We wanted to develop an open system that required minimal financial investment, assembly skills and computing knowledge. We settled on a design whose hardware is 100% mechanical, almost fully 3D-printable with PLA on low-end printers, assembled with glue, a pair of pliers and a cutter (Trinca and de Navascués 2026), and whose non-printable components can be bought from online retailers (**Table 1**). The accompanying software is minimal and includes step-by-step instructions for installation and self-guided notebooks to illustrate usage, aimed at users with no computing skills (Trinca and de Navascués 2026). The basic premise is, as in the Drosoflipper, simultaneously flipping several tubes, but with several differences. Instead of clipping the tubes at their opening, Drosben is based on racks that hold 12 tubes in honeycomb arrangement (**Figure 1A** and **Supplementary Figure 1A-C**), where flies are trapped by a collective lid (**Figure 1A** and **Supplementary Figure 1D-F**) that stays on robustly (**Supplementary Video 1**). Drosben racks are compact and can be laid on their base or their spine (**Figure 1B-C**), saving incubator space. Further, we took advantage of the asymmetric pattern of the tube array to create quick data recording sheets (**Supplementary Figure 2**). These can later be read using computer vision to collate the data automatically. To flip them, we use an intermediary chamber, the multiflipper (**Figure 1A** and **Supplementary Figure 1G-J**), so the position of the flies in tubes across the symmetry plane of the array stays the same. Flies are kept within the multiflipper chambers by a slider (**Figure 1A** and **Supplementary Figure 1K-M**), which closes all chambers at once (**Figure 1A** and **Supplementary Figure 1N-O**, compare with **Supplementary Figure G-I**). Drosben works in five steps, described next.

#### Registering the experimental conditions with the Drosben software

The details of the experimental design are captured in an Excel template, including the number of flies per each vial of a rack and their experimental variables (genotype, pharmacological treatment, sex, etc). The Drosben software will produce experiment-specific, printable data recording sheets (henceforth, ‘datasheets’), organised in series corresponding each to a specific rack (if the experiment uses more than one). It will also produce labels for the pouch in the spine of the rack. Datasheets and labels will match each rack with the experiment through a randomly generated alphanumeric ID and identify each rack within an experiment with a number; these can be made to bear explicitly the experimental conditions or made blind with a QR code (**Supplementary Figure 1P-Q**).

#### Loading flies in the initial Drosben rack

A rack with its corresponding label, loaded with fresh tubes (**Figure 1D**) and the depositor lid (**Supplementary Figure 1R-T**), is prepared beforehand. Animals anaesthetised with CO_2_ are counted, collected with a pooter and placed into each of the 12 fresh vials in the rack using the depositor rubber valves, and the depositor is replaced with a lid (**Figure 1D-G** and **Supplementary Video 1**). It takes less than 5 minutes to load a rack with 120 flies, which is below the estimated threshold of CO_2_ exposure before toxicological damage occurs (Linford, Bilgir et al. 2013, Shen, Yang et al. 2020).

#### Maintenance of lifespan experiments

Until the end of the experiment, flies will be periodically flipped into fresh food and the dead flies, usually stuck in the food of the old vials, recorded. This requires another rack, identified with the corresponding label and loaded with fresh tubes, and a multiflipper with the slider in (**Supplementary Figure 1N-O**). The lid of the old rack is substituted by the multiflipper, and each tube connected with one chamber of the multiflipper by removing the slider (**Figure 1H-I**). The multiflipper and the racked are flipped together, and flies are gently banged into the multiflipper chambers (**Figure 1J**). While the old rack is on top, it is exchanged for the new one (**Figure 1K-L**), and this is flipped again to place the multiflipper on top (**Figure 1M**) and bang the flies into the fresh vials—all in tubes with the same relative position within the array. Then, the multiflipper is substituted by the lid (**Figure 1M-N**), and the new rack can go back to the incubator (**Supplementary Video 2**). This will be repeated two or three days per week until all flies are either dead or a predetermined endpoint has been reached.

#### Data recording

Data recording takes place in the datasheet series corresponding to the rack being flipped. The correspondence is established in the header of the datasheet (**Supplementary Figure 3**) and the label of the rack, by the rack number and experimental details (or a QR code containing the experiment ID if the experiment is being carried out blindly; the code can be read by any smartphone camera) (**Supplementary Figure 1Q-R**). Datasheets contain grids of circles mimicking the array of tubes within a rack (**Figure 1O-P**). Each grid corresponds to an observation and flipping date; each event is recorded simply by making a mark on the circle of the grid with the same relative position as the tube where the event was detected in the rack (**Figure 1P**). In the simplest scenario, the events are deaths, detected as immobile flies stuck at the bottom of the old tube. However, dead flies may fall to the new tube or stuck flies may still be alive; alive flies may escape. To account for this, Drosben allows three different event types: death, censoring (escaped, stuck alive) and ‘carried-overs’ (dead but not stuck). Each is identified with a different colour, so the user only needs to record what they see with the corresponding marker pen; the software will identify the event type by the colour, back-date the carried-overs when they finally get stuck to the food, and keep a tally of total events to match the original number of flies. Before the datasheets are generated, the software checks that the colours produced by the user’s pen markers can be distinguished automatically. We have pre-tested multiple combinations that work well, including some that are colourblind-friendly (**Figure 2A**).

**Figure 2.**
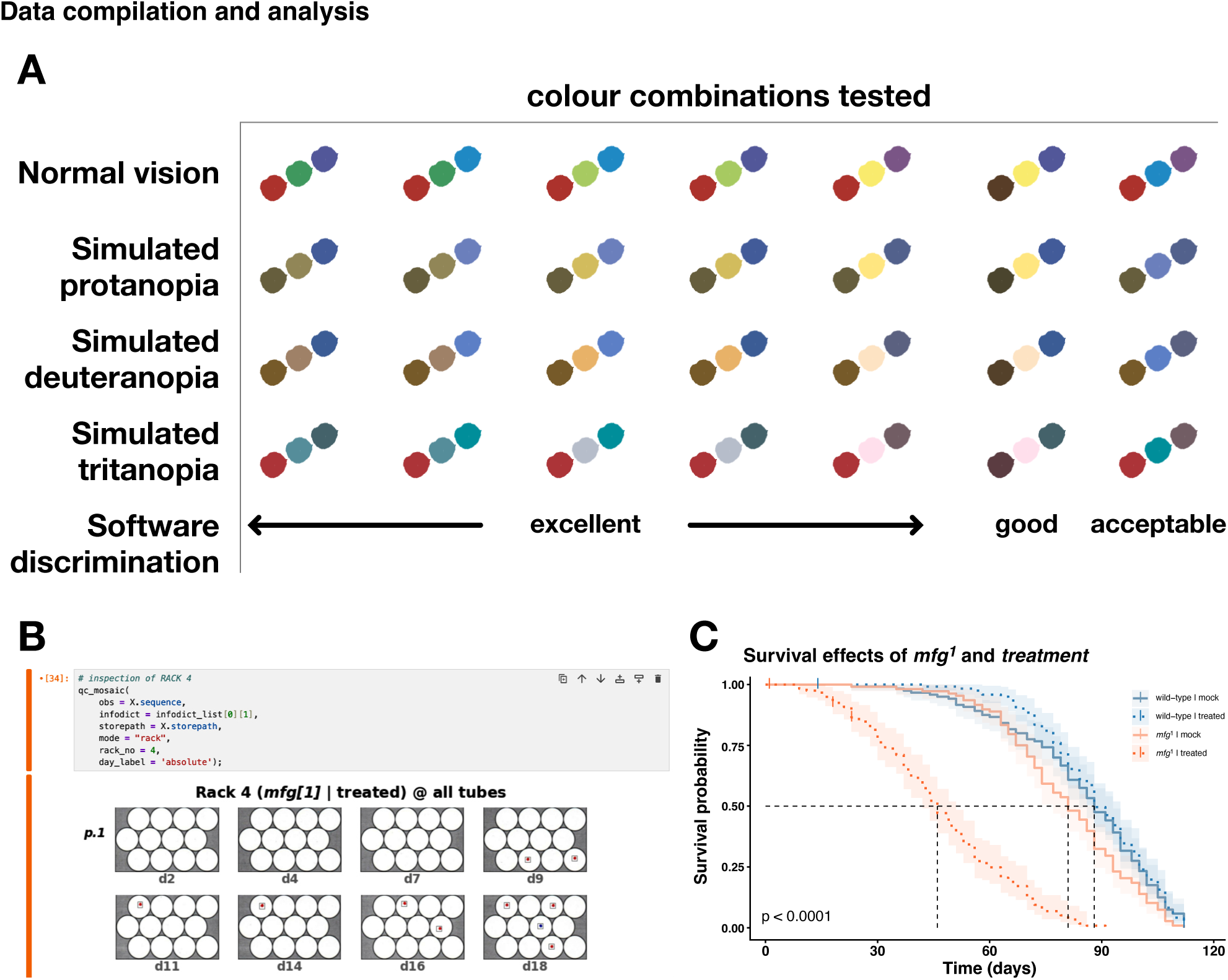
Drosben allows flexible recording of data. (**A**) Combinations of three coloured markers used to record separately deaths, carried-overs and censorings in the datasheets. The colours are a close approximation, as the exact colourmap values depend on the marker ink, the paper used and the scanning device. Simulated colourblind was made using DaltonLens (Burrus 2025) based on the colour models described in (Brettel, Vienot et al. 1997). The main discriminatory value used by the Drosben software is hue, so colourblind-friendly combinations reduce the available colourmap range and may not perform as well, but they can be used. (**B-C**) Data compilation and analysis. Scanned datasheets are analysed with the Drosben software running in interactive notebooks (**B**) that also allow the inspection of the recognised events for manual corrections. Life table files produced by the notebooks are used directly in R scripts to produce Kaplan-Meier plots (**C**) and bespoke statistical tests. Graph in C depicts the synthetic data included in the software installation for testing and training.

The mechanics at the bench to identify the events is as follows. First, the old rack is inspected to record the immobile flies stuck to the food as deaths. Stuck flies that are still moving are recorded as censored. Also censored are any flies that escape. If the tube these escaped from is not known, they can be recorded in any tube; at the end of the experiment, the software will re-assign censorings from tubes with excess events to tubes missing events. Then the new rack is inspected for immobile flies at the bottom of tubes, that remain immobile after a few flicks; these are recorded as carried-overs. With 95 mm-tall tubes, the the bottom of the tubes in internal positions of the rack is visible at an angle; with shorter ones, the view may be blocked by the lid. In this case, carried-overs should be recorded earlier, before finishing the flip, while the flies are in the multiflipper chambers, and carried-overs fall on the mesh or the slider, depending on the orientation of the multiflipper.

If the user realises that they made recording mistakes (e.g. marking on the wrong place or with the wrong colour), correction fluid or tape could be used to redress this. If the mistake affects many marks, or the datasheet has been defaced by stain or crumpling, the easiest thing is to make a note with pen or pencil that explains the necessary correction and continue with passaging. Later, the user can reprint the specific (rack and page) datasheets with mistakes and correct them. Re-populating a blank datasheet with marks from another is trivial and takes seconds. Some non-advertent mistakes will be identified by the software, which will take a tally of the observed deaths and censorings and compare with the declared number of flies per tube, warn the user of mismatches and propose a correction course.

#### Data logging and analysis

At any point in the experiment, the user can scan the recorded datasheets to compile the data and generate a spreadsheet with the life table up to that time. If the experiment has not finished, flies alive at that point will be right-censored, allowing the user to follow the progression of the experiment. The datasheet scans will be ‘read’ individually by the Drosben software, which will assign each scan to an experiment and a rack within that experiment, so multiple experiments can be digitalised at once without the need to organise the scan files. The interaction with the user is through Jupyter notebooks (**Figure 2B**), which allow performing some quality control of the event recognition and deciding how to censor the survivor flies. Once the life tables have been generated, users can analyse them using their own scripts or use the R markdown notebook provided to obtain a Kaplan-Meier curve (**Figure 2C**) as well as estimates of the magnitude of effect of variables and their interactions in the form of hazard ratios, using a basic Cox regression model.

### Drosben reduces handling time without affecting *Drosophila* lifespan

Having established the system, we wanted to test whether it was usable in practice or had a deleterious effect on survival that could be confounding. To do this, we asked a novice (three months of experience handling *Drosophila* vials) and an intermediate (over 1 year of experience) user to perform parallel survival experiments using Drosben and conventional handling. Both users managed to finish the lifespan experiment without major setbacks (e.g. excessive censoring). In the hands of the novice user, the Drosben hardware had a negative effect on survival, as indicated by Cox regression (HR 1.4, *p*=0.04) (**Figure 3A**). However, this was not the case with the intermediate user, whose use of Drosben was protective (HR=0.8), though this was insignificantly (*p*=0.07) (**Figure 3C**). When observing the novice, it was clear that they banged the flies with excessive force when transferring them between multiflipper and rack. Flies are sensitive to mechanical harm, which leads to lifespan reduction that worsens with increased banging (Katzenberger, Loewen et al. 2013, Barekat, Gonzalez et al. 2016). Therefore, we attribute the reduced lifespan when Drosben was used by the novice experimenter to accumulative mechanical trauma derived from a sub-optimal handling technique. From our results we conclude that using Drosben does not intrinsically bring a survival handicap and will capture lifespan changes when all experimental arms are handled with it.

**Figure 3.**
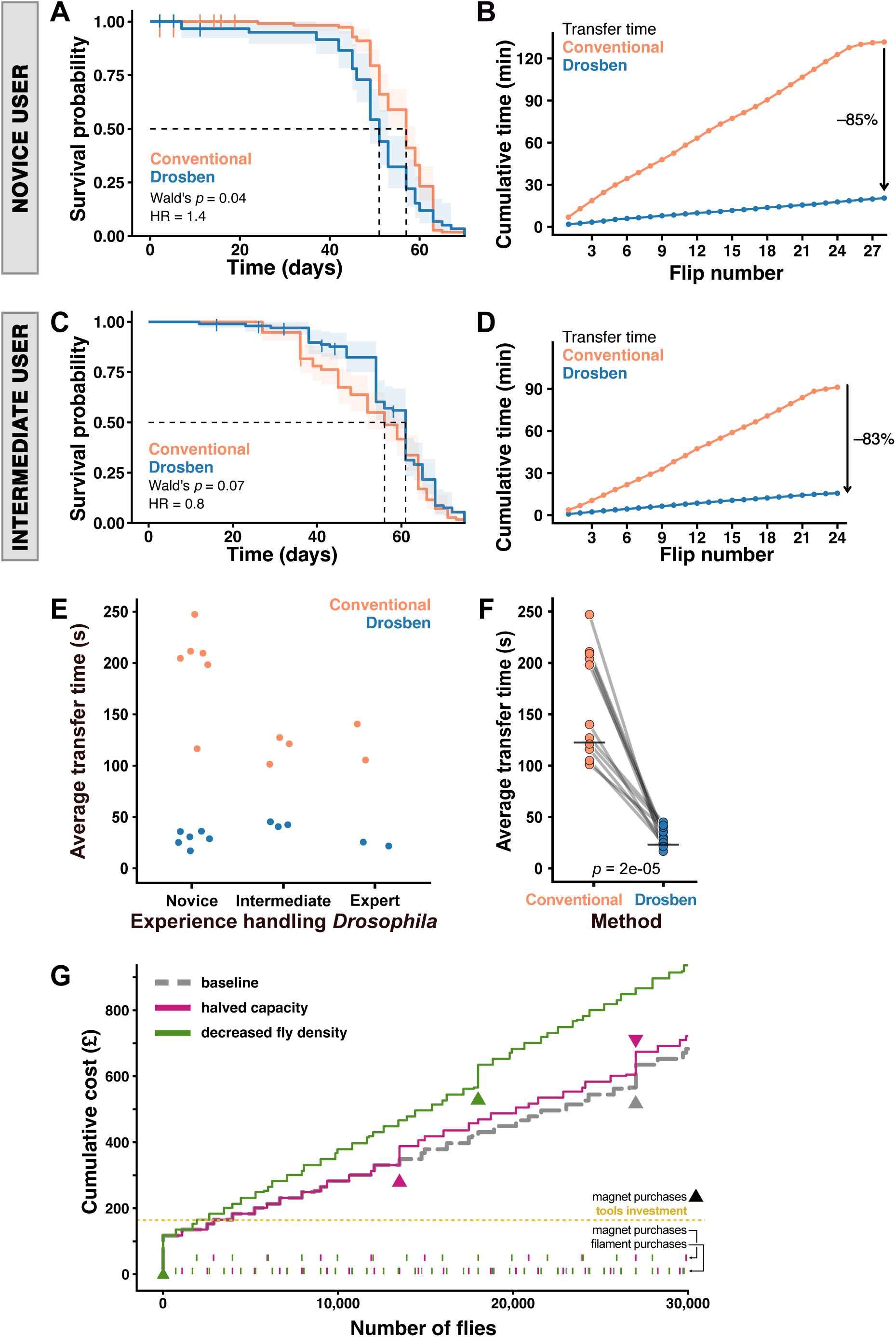
Drosben improves handling time across experience levels. (**A-D**) Details of Drosben usage results by one novice and one intermediate user. (**A, C**) Kaplan-Meier plots with handling method as covariate for a novice (**A**) and intermediate user (**C**). Each curve corresponds to a cohort of 120 female *w^1118^* flies housed in 12 tubes until all were dead or censored (vertical strokes). *P* values correspond to Wald tests from Cox regression. HR, hazard ratio. (**B, D**) Cumulative transfer time according to handling method for the experiments depicted in **A** and **C**. (**E-F**) Comparison of transfer speed of volunteers, by handling method. Transferred cohorts were 120 *w^1118^* females. The recorded times were the cumulative time taken for 12 consecutive transfers (for conventional) and the average of 3 consecutive transfers (for Drosben). (**E**) Transfer times according to experience level with *Drosophila* husbandry (novice: < 4 months; intermediate: 4–18 months; expert: > 18 months). (**F**) Overall comparison of transfer time (paired t-test). (**G**) Cost behaviour of the Drosben hardware with workload expressed as the total number of flies handled in parallel. Costs were calculated from prices in Table 1. The ‘baseline’ curve (grey) assumes: (1) 15 flies will be housed per tube, (2) handling takes 32 seconds per rack (the average flipping time, with all experience levels weighted equally), and (3) the experimenter can commit 4 hours of flipping time per passage day (so, 12 per week); this is continuous flipping time, with recording, preparing food or going back and forth from the incubator not accounted for (hence the limit of 4 hours). The curve displaying ‘halved capacity’ (magenta) shows cost behaviour when flipping time is reduced to 2 hours per passage or the user is slower (64 seconds per flip, which we have never recorded). The curve displaying ‘decreased density’ (green) shows cost behaviour when tubes contain 10 flies per tube (note the number of cohorts or experiments could be the same, but with lower statistical power). Coloured triangles show when the cost is brought up by the fabrication of a new multiflipper and slider (mostly driven by the acrylic tubes). Yellow line shows the additional cost of starting without any tools (3D-printer, pliers, cutter and silicone sheet). The rows of coloured ticks at the bottom show when the cost is brought up by the purchase of a PLA filament roll or a pack of magnets, as indicated, in the corresponding colour curve.

We next wanted to confirm that Drosben reduced the handling time of survival experiments. We recorded the time taken for a novice and intermediate users to perform parallel lifespan measurements with both transfer methods. In both cases, the Drosben method reduced time invested ∼6 fold: during the course of the lifespan experiment, the novice user spent 130 minutes handling flies conventionally, but only 20 minutes using Drosben (85% reduction; **Figure 3B**); the intermediate user invested 91 minutes flipping vials conventionally versus 16 minutes using Drosben (83% reduction; **Figure 3D**). To expand these observations to researchers who were not involved in the design of the Drosben hardware, we recruited 11 colleagues with experience in *Drosophila* handling ranging from 4 months to 20 years. We grouped them as novice (<4 months), intermediate (4-18 months) or expert (>18 months of experience), and asked them to transfer 12 tubes both conventionally and using Drosben. As expected, more experienced Drosophilists were faster at transferring conventionally than novices, and as a result novice users benefitted the most from using Drosben (**Figure 3E**). Despite this, all users benefit from shorter handling time (∼3– to 5-fold) when using Drosben (**Figure 3F** and **Supplementary Video 2**).

### Drosben costs scale with the amount of generated data

Our aim was for Drosben to save both time and money, so we used cost behaviour analysis to determine whether the costs of fabrication would undermine the benefits in handling speed. We realised that the increasing costs of having to handle more experiments running simultaneously scales slowly with the number of flies: in our ‘baseline’ conditions (committing four hours of continuous flipping three times a week, flipping a rack in 32 seconds and housing 15 flies per tube), after an initial investment of ∼£120 (∼£300 if starting without any tools), one needs to plan to process over 10,000 flies at a time before having to spend another ∼£120 in fabrication materials (**Figure 3G**). In these conditions, a single user with one multiflipper could handle up to ∼27,000 flies; beyond that number, a second user and multiflipper would need to be engaged. We evaluated the effects of halving the work capacity (reducing the time committed to flipping to 2 hours per passaging day or increasing the flipping time to 64 seconds per rack); logically this brought forward the need to engage a second user and multiflipper to the point of handling over ∼13,500 flies (**Figure 3G**, grey and magenta triangles), but the overall costs are not significantly increased. By contrast, reducing the housing density from 15 to 10 flies (**Figure 3G**, green curve) did increase the slope of the cost behaviour, as expected: new racks and their lids need to be printed with a smaller number of total flies. In any case, the cost behaviour shows that a laboratory can handle up to 30,000 flies in parallel for a one-time cost of ∼£900.

### Static magnetic fields up to the milli Tesla range do not reduce *Drosophila* lifespan

We added rare-earth magnets to the design of Drosben racks and lids to secure the latter without a reduction in handling speed. It has been reported that *Drosophila* can sense magnetic fields which can impact fitness through impairing neurological function, including short-term survival to starvation (Gegear, Casselman et al. 2008, Fedele, Green et al. 2014, Kawasaki, Okano et al. 2023). As the magnetic field generated by a magnet decreases sharply with the distance to it, the design of the Drosben multiflipper provided the opportunity to test the effect of magnetic field on longevity with a range of field intensities. We obtained estimates for the magnetic field generated by the magnet at different heights along the central axis of each tube; this shows that, in the closest tubes to the magnet, flies could be repeatedly experiencing magnetic fields in the mT range (**Figure 4A** and **Table 2**), at least tens of times the range of intensity of the Earth’s magnetic field (25-65 µT) (Hulot, Finlay et al. 2010). However, *Drosophila* lifespan did not depend on the vial proximity to the magnet (**Figure 4B**). From this, we conclude that *Drosophila* can tolerate exposure to magnetic fields considerably larger than the Earth’s.

**Figure 4.**
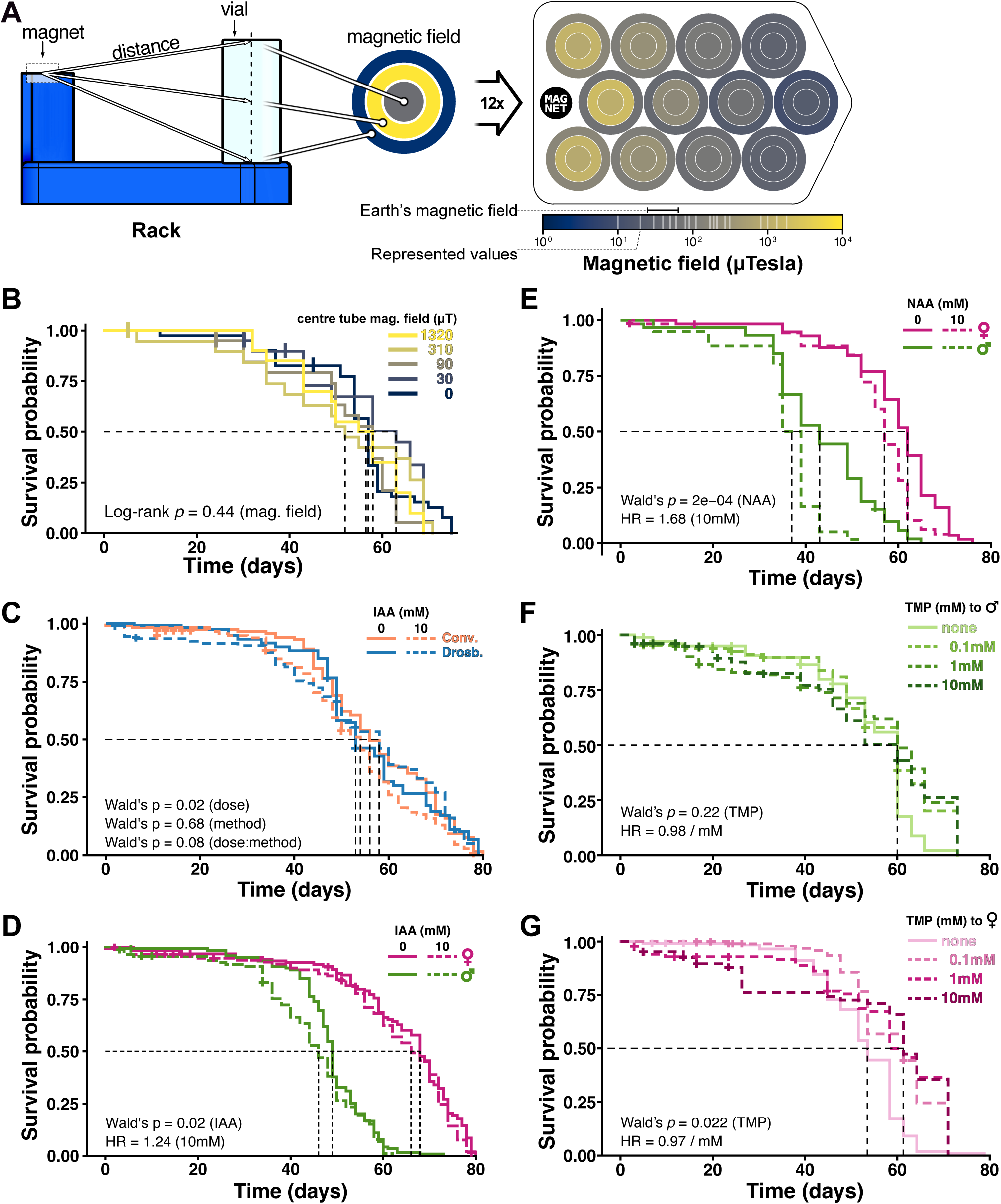
Effects on lifespan of magnetic fields, auxins and trimethoprim. (**A**) Left: schematic depicting the points considered for estimation of the magnetic fields. We calculated the coordinates from the centre of the magnet (two touching N52 Neodymium-Iron-Boron discs, 12 x 3 mm each) to the top, middle and bottom points of the central axis of each food vial. Right: heatmap of estimated magnetic fields values (see Table 1) at the three heights of each tube, as indicated. Note the colourmap is in logarithmic scale and shows the range of Earth’s magnetic field strength and the values represented in the heatmap. (**B**) Kaplan-Meier plot for *Vallecas* males with distance to magnet as covariate (where infinite is a negative control with no magnet). N=20 flies per curve (except for the control, where N=40). Note that the colourmap values are different from A. The *p* value is from a Log-rank test. (**C, D**) Kaplan-Meier plots for *Vallecas* flies treated with 10 mM IAA added to the food throughout adult life, transferred conventionally or with Drosben. Covariates considered were handling method, treatment and their interaction (**C**), or treatment and sex (**D**). N=120 flies per curve. *P* values are from Wald tests for individual covariates in Cox regression. (**E**) Kaplan-Meier plot for *Vallecas* flies treated with 10 mM NAA added to the food throughout adult life, transferred with Drosben, representing treatment and sex as covariates. N=60 flies per curve. The *p* value is from a Wald test for Cox regression, with only treatment as covariate. (**F, G**) Kaplan-Meier plots for *Vallecas* males (**F**) and females (**G**) treated with 0.1, 1 and 10 mM TMP in the food throughout adult life, transferred with Drosben. N=60 flies per curve. *P* values are from Wald tests for Cox regression, with dose as continuous covariate. For all panels, flies were housed ten per vial until all were dead or censored (vertical strokes). HR: hazard ratio.

**Table 2.** Estimated magnetic field intensities in the racked tubes. Tube numbering proceeds in rows first, then columns, from top to bottom and left to right, considering the tube grid as oriented in Figure 4A. Values of magnetic field were estimated using the KJ Magnetics Ltd (Pennsylvania, USA) online field calculator (K&J Magnetics Inc. 2025), for a rare-earth magnet grade N52, 6 mm diameter and 6 mm thickness (for two 3 mm thick magnets, one each in the rack and the lid), at the distances between the magnet and the bottom, middle and top of the central axis of each tube. Values in blue are within the range of the of the Earth’s magnetic field.

| tube | Magnetic field ($\mu\text{T}$) | | |
| --- | --- | --- | --- |
|  | bottom | middle | top |
| 1 | 170 | 1320 | 810 |
| 2 | 110 | 310 | 270 |
| 3 | 50 | 90 | 80 |
| 4 | 30 | 30 | 30 |
| 5 | 60 | 1790 | 1000 |
| 6 | 40 | 210 | 190 |
| 7 | 20 | 60 | 60 |
| 8 | 10 | 20 | 20 |
| 9 | 170 | 1320 | 810 |
| 10 | 110 | 310 | 270 |
| 11 | 50 | 90 | 80 |
| 12 | 30 | 30 | 30 |

### Auxins mildly decrease lifespan while trimethoprim expands female lifespan

The repertoire of ligand-regulated degron technologies is growing quickly, as the ability to exert acute depletion of proteins allows much better insight into protein function in very dynamic processes and has many potential applications in technology and medicine (Kanemaki 2022, Shi, Zhao et al. 2025, Ma, Yin et al. 2026). In *Drosophila*, the AID and DD degron systems have been employed for specific genes or transgenes (Trost, Blattner et al. 2016, Bence, Jankovics et al. 2017, Chen, Werdann et al. 2018, Kogenaru and Isalan 2018, Lopez Del Amo, Leger et al. 2020, Jullien, Guillou et al. 2022) and also incorporated in general-purpose transgenes to add a layer of drug control to the Gal4/UAS system for targeted misexpression (Sethi and Wang 2017, McClure, Hassan et al. 2022). Therefore, we sought to use Drosben to test the effect of the ligands for these systems (IAA and NAA for AID, TMP for DD) at concentrations used to regulate the degrons (up to 10 mM for each compound). First, we tested IAA with both conventional handling and Drosben. Using Cox regression, we found that treatment with IAA at 10 mM had a small but significant effect (HR_IAA_=1.37), but the handling method did not, nor this had a significant interaction with treatment (**Figure 4C**). As mifepristone/RU486, another drug used to control transgene expression, has a (mated) female-specific effect on lifespan (Ford, Hoe et al. 2007, Landis, Salomon et al. 2015) we tested whether this was the case for IAA, and found no significant interaction between sex and treatment as variables (**Figure 4D**). Using only Drosben, we next tested NAA and TMP. NAA at 10mM had stronger effects than IAA (HR_NAA_=1.68), which were also non-sex-specific (**Figure 4E**). We tested TMP at a range of concentrations and found that it only significantly affected the lifespan of females, for whom it was protective (HR_TMP_ (per mM) = 0.97; HR_TMP_ (for 10 mM) = 0.73) (**Figure 4F-G**). From this, we conclude that chronic exposure as adults to 10 mM of any of the three degron-regulating ligands have noticeable effects on lifespan.

## DISCUSSION

Measuring lifespan is common in multiple *Drosophila* research fields and labour-intensive. Here we describe the Drosben system for survival analysis of *Drosophila*, which we developed for adoption with minimal financial or upskilling investment (**Figure 1**, **Supplementary Figure 1** and **Table 1**). We have shown that this system reduces handling time 3– to 5-fold, with benefits regardless of *Drosophila* husbandry experience but higher benefits for novice users (**Figure 3**). We have also demonstrated the capacity to generate high-quality lifespan datasets (**Figure 4**). We expect using Drosben would help scaling-up lifespan experiments to power larger or more complex experiments; for instance, genome-wide association studies (GWAS) of *Drosophila* lifespan have had to use cohort sizes ranging from 21 to 72 flies per genotype (Durham, Magwire et al. 2014, Ivanov, Escott-Price et al. 2015, Huang, Campbell et al. 2020), which are relatively small compared to what an individual *Drosophila* lifespan experiment would typically employ.

The design of the Drosben components could be easily adapted for food vials with different dimensions, or to extend and improve its capabilities. We have only tried this transfer method with polystyrene vials, but not with glass ones; these might require padding the rim of the multiflipper chambers with rubber. Also, the deleterious effects of using Drosben in the hands of a novice prompted us to introduce the Drosben to new users with a single ∼30 minute ‘dry run’ to transfer flies between multiflipper and racked vials. However, banging could be halved, and speed improved, by directly transferring the flies from the old rack to the new one; we chose to avoid this because flies would change tube positions across the symmetry plane of the rack every flip, and we thought it could make tracking the behaviour of individual tubes more difficult. The same transfer logic could be applied to larger containers for cohorts of thousands of individuals, as would be needed to estimate mortality rates.

Including magnets in the design of Drosben to improve its speed made us consider the effect of magnetic fields on *Drosophila* lifespan. It has been reported that *Drosophila* can sense magnetic fields, and these influence their negative geotaxis behaviour, survival to starvation, or circadian clock, among other traits (Gegear, Casselman et al. 2008, Yoshii, Ahmad et al. 2009, Fedele, Green et al. 2014, Kawasaki, Okano et al. 2023). More recently however, it has been claimed that there is no evidence of magnetic fields influencing *Drosophila* behaviour (Bassetto, Reichl et al. 2023). We found no evidence that the magnetic fields produced by the magnets in the Drosben rack and lid modulate lifespan (**Figure 4B**), even though some areas of the tubes would have field strengths in the mT range (**Figure 4A** and **Table 1**). However, we cannot rule out that some strains or mutants may be sensitive to magnetic fields of the strengths generated by the magnets employed in the Drosben system.

Finally, we described the effect on lifespan when *Drosophila* are chronically exposed to three compounds used in targeted protein degradation: indole-3-acetic acid (IAA), 1-naphthalene-acetic acid (NAA) and trimethoprim (TPM). Both auxins reduce *Drosophila* longevity, with NAA having a stronger effect than IAA. TMP extends the lifespan of females. The effect of TMP on *Drosophila* life history traits has not been evaluated before, but our observation contrasts with its reported effect on *Anopheles stephensi*, where exposure to concentrations of 1-10 mM show strong deleterious effects in larval and adult survival (Zamy, Futo et al. 2026). The effect of auxins on *Drosophila* physiology has been documented before, out of their interest as a degron regulator in heterologous expression systems as well as the importance of indoles as microbiome-derived catabolites of dietary tryptophan and agonists of the aryl hydrocarbon receptor (AHR) (Hubbard, Murray et al. 2015). McClure et al. (2022), when describing their *AGES* method for the temporal control of transgene misexpression, show that exposure to 5-10 mM of NAA do not affect survival of larvae nor adults, though 10 mM delays development by a day (McClure, Hassan et al. 2022). Other authors have reported deleterious effects of both NAA (Fleck, Biswas et al. 2024) and IAA (Aziz, Ramesh et al. 2024), but in both cases at concentrations higher than 10 mM (10-350 mM of NAA and 60 mM of IAA). However, Fleck et al., (2024) detect other effects at 10 mM, such as developmental delay, changes in adult feeding behaviour (less frequent, shorter interactions with food) and lower triglyceride levels. By contrast, different paradigms of indole exposure report beneficial effects. Indole-3-propionic acid, another naturally occurring auxin that is also produced in the dietary tryptophan catabolism by human gut microbes, seems to mildly expand *Drosophila* longevity as a dietary supplement at ∼16 mM; the effect is modulated by sex and genetic background (Vyavahare, Berger et al. 2026). Using mono-association of *Drosophila* with wild-type and Tryptophanase mutant strains of *Escherichia coli*, Sonowal et al. (2017) showed that bacterial IAA prolonged adult lifespan through the activity of the AHR, since zygotic mutants for *spineless* (which encodes *Drosophila* AHR) did not show this benefit (Sonowal, Swimm et al. 2017). These observations are difficult to reconcile, as the specific auxin, concentration, form of exposure, genotype, diet, and associated microbiome in each study are not comparable. However, these studies as well as our own work strongly suggest that auxins are not orthogonal molecules to *Drosophila* physiology, and we concur with (Fleck, Biswas et al. 2024) in that this should be considered when using auxin-induced degrons in experimental designs. This probably applies to TMP-regulated degrons too (**Figure 4F-G**).

In summary, we have developed the Drosben system to streamline the labour-intensive lifespan assays in *Drosophila* and demonstrated its capacities. We described it in full, in the spirit of open hardware, providing a stable guidance for usage and manufacturing, to encourage adoption and adaptation (Trinca and de Navascués 2026). The design aims to be affordable for any research group irrespective of funding level, manufacturing capacity or computing skills, to support *Drosophila* research in a wide range of areas of direct biomedical interest.

## AUTHOR CONTRIBUTIONS

TMT: Data curation, formal analysis, investigation, methodology, software, supervision, writing – original draft, writing – review & editing. PBM and CFG: Investigation, validation, writing – review & editing. JdN: Conceptualization, data curation, formal analysis, funding acquisition, methodology, software, supervision, writing – review & editing.

## Supporting information

Supplemental File 1 with source data

## ACKNOWLEDGMENTS

We thank the *Drosophila* researchers who volunteered to compare the handling speed using Drosben and conventional flipping, and our *Drosophila* colleagues at the School of Biosciences of Cardiff University for useful discussions and suggestions while developing the system.

## SUPPLEMENTARY FIGURE LEGENDS

**Supplementary Figure 1:**
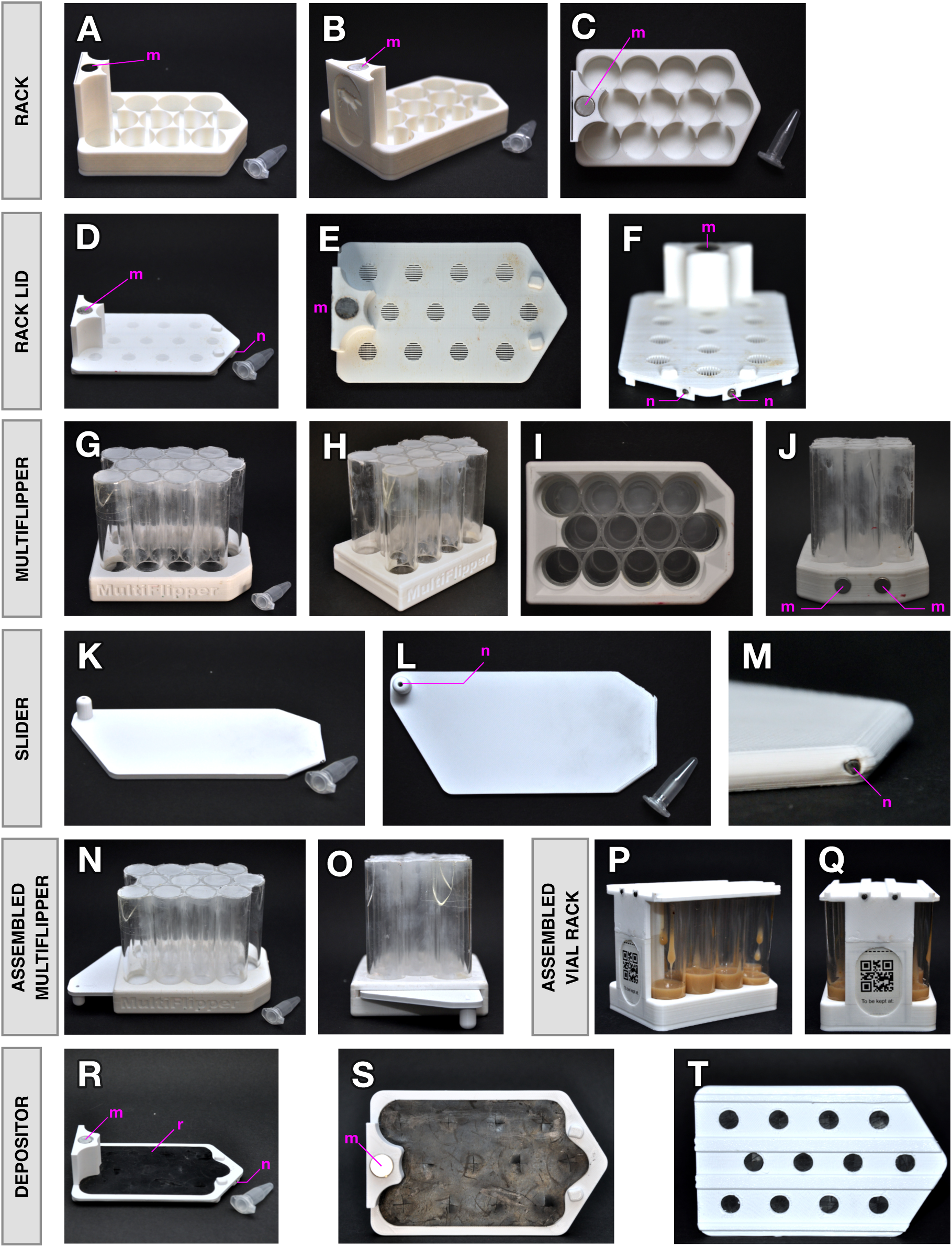
Individual Drosben components. Photographic description of the five 3D-printed components of the Drosben hardware: rack, its lid, multiflipper, its slider, and depositor. Throughout the figure, ‘m’ indicates a magnet, ‘n’ a nail and ‘r’ a rubber sheet glued to the part. A microcentrifuge tube is placed in many panels as size reference. (**A-C**) Top-side, back-corner and top views of the rack to house the food vials. Note in **B**, **C** the sleeve in the spine of the rack to hold a printed label (see panels Q, R). (**D-F**) Top-side, top and top-front views of the lid. Note the breathing slits and the two catch buttons (**E**, **F**). (**G-J**) Top-side, back-corner, bottom and front views of the multiflipper. Note the glued (nylon) mesh (**G**, **H**), the magnets (**J**) and the base of the acrylic tubes, glued to the top side of the multiflipper structure, flush with the slit for the slider (**I**). (**K-M**) Top-side, top and detail views of the slider. Note the opening for a nail (optional) on the slider puller (**L**). The nail in **M** is required for the multiflipper magnets in **J** to pull and secure the slider. (**N-P**) Top-side, top and bottom views of the depositor. Note the cuspid valves remain closed when not crossed by the pooter as in Figure 1E. (**Q-R**) Back-corner and back view of the rack containing 12 tubes covered by the lid. Note the label inserted in the rack spine sleeve. (**S-T**) Top-side and back views of the multiflipper with slider. Note the slider closes the acrylic tube chambers.

**Supplementary Figure 2.**
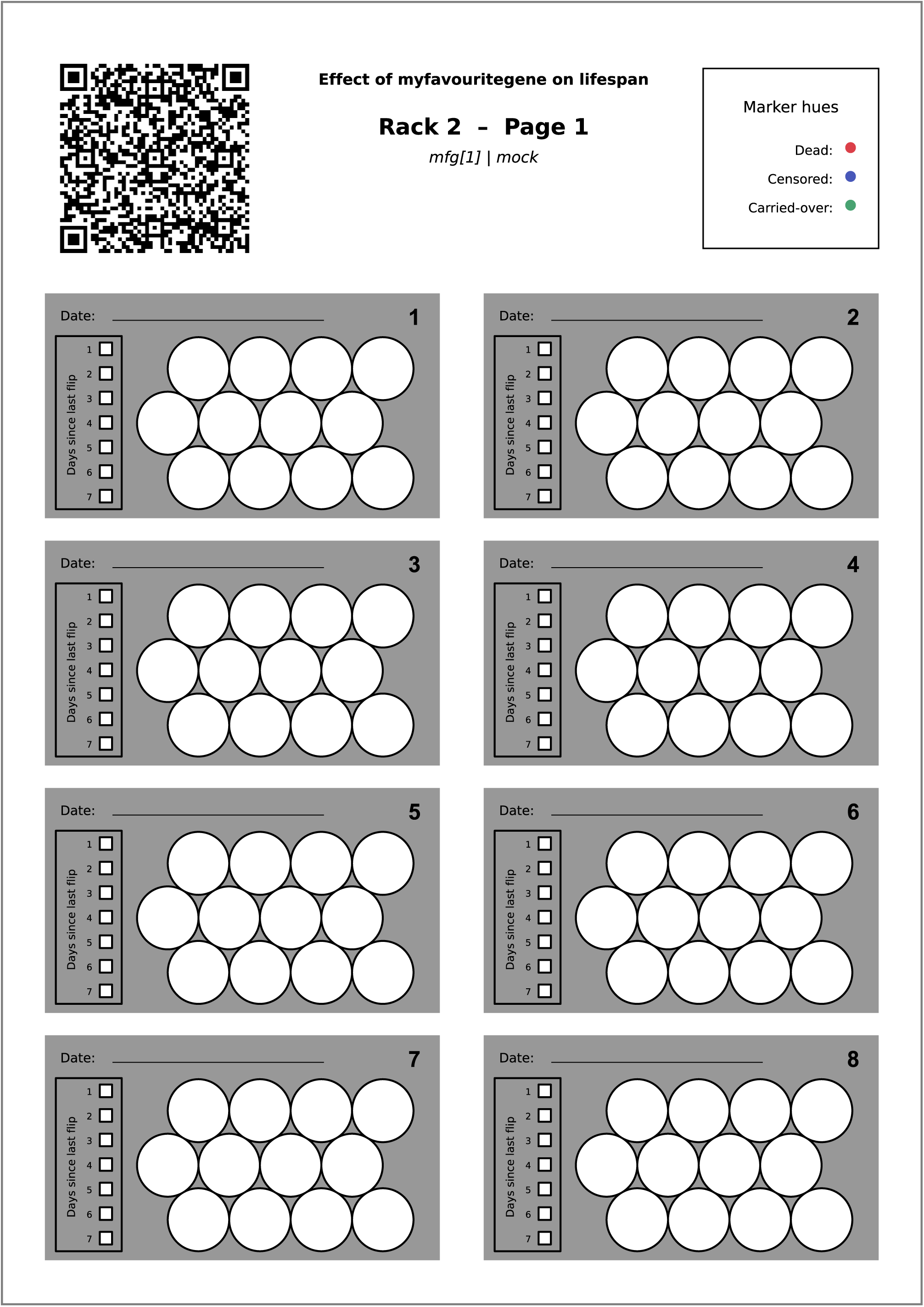
Organisation of the Drosben datasheets. Data recording sheet of a fictional experiment (non-blind version). The header identifies the experiment, the rack, the datasheet page, the experimental arms contained in that rack and the expected correspondence between colours and event types. The QR code is used similarly by the computer vision pipeline of the software. Below the header, the blocks of circle arrays represent the top view of the old tubes in the rack, to be used in eight observations chronologically ordered. Each block contains a line of squares to mark the days between that observation and the previous.

## SUPPLEMENTARY MOVIE LEGENDS

**Supplementary Movie 1. The Drosben method for lifespan assaying and analysis.**

Related to Figure 1. The rack lid stays in place when the rack is knocked over in multiple directions. A rack dropped from a height will not keep the lid on, but simply by adding a rubber band (size 64) secures them against almost any drop.

**Supplementary Movie 2. The Drosben method for lifespan assaying and analysis.**

Related to Figure 1. Demonstrations using the Drosben hardware. Step 1: Initiating an experiment using the rack, depositor (with a pooter) and lid to load flies into vials. Step 2: flipping survivor flies into fresh vials using a second rack, the multiflipper with slider, and the lid.

**Supplementary Movie 3. Conventional flipping versus the Drosben method.**

Related to Figure 3. An experienced user flipped 12 vials and recorded deaths using the conventional and Drosben handling methods, shown in parallel to demonstrate the transfer and recording speed benefits of Drosben.

## SUPPLEMENTARY FILES LEGENDS

### Supplementary File 1. Data for all figures

XLSX file containing the data for all figures, each in one spreadsheet in tidy format.

